# Community size affects the signals of ecological drift and niche selection on biodiversity

**DOI:** 10.1101/515098

**Authors:** Tadeu Siqueira, Victor S. Saito, Luis M. Bini, Adriano S. Melo, Danielle K. Petsch, Victor L. Landeiro, Kimmo T. Tolonen, Jenny Jyrkänkallio-Mikkola, Janne Soininen, Jani Heino

## Abstract

Ecological drift can override the effects of deterministic niche selection on small populations and drive the assembly of small communities. We tested the hypothesis that smaller local communities are more dissimilar among each other because of ecological drift than larger communities, which are mainly structured by niche selection. We used a unique, comprehensive dataset on insect communities sampled identically in a total of 200 streams in climatically different regions (Brazil and Finland) that differ in community size by fivefold. Null models allowed us to estimate the magnitude to which beta diversity deviates from the expectation under a random assembly process while taking differences in species richness and relative abundance into account, i.e., beta deviation. Beta diversity of small tropical communities was consistently higher but closer to null expectations than β-diversity of large communities. However, although β-deviation and community size were strongly related in both regions, the direction of the relationship varied according to dissimilarity metrics. While incidence-based β-diversity was lower than expected (communities were less dissimilar than null expectations) and negatively related to community size in Brazil, abundance-based β-diversity was higher than expected (communities were more dissimilar than null expectations) and positively related to community size in both regions. We suggest that ecological drift drives variation in small communities by increasing the chances of species with low abundance and narrow distribution to occur within the metacommunity. Also, while weak niche selection and high dispersal rates likely reduced variation in community structure among large tropical streams, niche selection was likely sufficient to cause non-random variations in the relative abundances of genera among large communities in both regions. Habitat destruction, overexploitation, pollution, and reductions in connectivity have been reducing the size of biological communities. These environmental pressures will make smaller communities more vulnerable to novel conditions and render community dynamics more unpredictable, as random demographic processes should prevail under these conditions. Incorporation of community size into ecological models should provide conceptual, empirical and applied insights into a better understanding of the processes driving changes in biodiversity.

## Introduction

Recent syntheses in community ecology propose that four main processes drive the dynamics of metacommunities – deterministic niche selection, ecological drift, dispersal and speciation (Vellend 2010, 2016, Leibold and Chase 2018). At broad spatial scales, dispersal, colonization history, and spatial heterogeneity are thought to produce major biodiversity patterns (Leibold and Chase 2018), while speciation also plays a role by altering the composition of regional species pools at longer time frames (Vellend 2010). Within localities, niche selection determines community structure mainly through species interactions, differential use of resources and species responses to environmental gradients (Leibold and Chase 2018). However, stochasticity may also play a major role in driving local community dynamics, for example, when demographic events occur at random with respect to species identities (Vellend et al. 2014). Indeed, mechanistic models (Orrock and Fletcher 2005, Orrock and Watling 2010) and recent empirical evidence (Gilbert and Levine 2017) suggest that ecological drift can even override the effects of niche selection under certain circumstances, such as when species populations in local communities are small and isolated from other populations.

Small communities have few individuals per unit area, and thus random birth and death events are likely to have a high impact on their structure (species composition and relative abundances). For example, the species composition of a local community would change if all individuals of a species die before reproducing. This is likely to happen in nature especially among small populations or in those large populations in which only a reduced fraction of adults successfully reproduce (Bunn and Hughes 1997). Some models indicate that ecological drift can even reduce competition asymmetries in small communities to a level that strong and weak competitors become effectively neutral (Orrock and Watling 2010) – i.e., the negative effect of a superior competitor on other species is relatively small compared to the effects of demographic stochasticity. Indeed, in an experiment with annual plants, Gilbert and Levine (2017) found that larger communities converged to a state in which a strong competitor dominated after three years, whereas in smaller communities the strong competitor co-occurred with other species at different densities. Thus, both theory and experimental evidence suggest that ecological drift can change the structure of local communities by altering species relative abundances as well as species occurrences. Investigating the relationship between variation in community structure and community size would help us better understanding the relative importance of deterministic and stochastic processes on beta diversity. This should also be relevant from an applied perspective, as many types of environmental disturbances, such as deforestation and floods, tend to reduce the size of local communities (Barnes et al. 2014, Petsch et al. 2015).

Most methods commonly used to estimate β-diversity are affected by differences in species richness or species abundance distributions (Chase and Myers 2011). This is undesirable for understanding the beta diversity-community size relationship for two main reasons. First, estimates of β-diversity can be influenced by random sampling effects that are neutral with respect to species identity (Chase and Myers 2011, Myers et al. 2013, 2015). For example, let us assume that the species composition of 10 local communities embedded in a larger metacommunity represents each a random subset (say 5 species) of the regional species pool with, for instance, 100 species. This would likely result in high β-diversity as, by chance, many pairs of local communities would not share species. Second, studies employing incidence-based estimates of β-diversity (e.g., Jaccard or Sørensen dissimilarity indices) do not capture variation in species relative abundances (Anderson et al. 2011). In this case, if the same species occur at two sites but with different relative abundance, β-diversity should not be equal. A solution to the first issue is to use a null model to produce expected values under random assembly from a large species pool, contrast observed and expected values and use the difference between them as estimates of β-deviation; i.e., the extent to which β-diversity differs from null expectations (Kraft et al. 2011, Myers et al. 2013, 2015, Catano et al. 2017). In this case, positive and negative values of β-deviation indicate that communities are more dissimilar and less dissimilar than expected by chance, respectively. Beta deviation values close to zero indicate communities are as dissimilar as expected by chance (Kraft et al. 2011, Chase et al. 2011, Catano et al. 2017). Beta deviation provides a means to determine whether β-diversity deviates from the expectations of stochastic assembly and if the magnitude of the deviation is related to community size. A solution to the second issue is to analyze the data with dissimilarity coefficients that take into account species composition and species relative abundance (e.g., Bray-Curtis; Anderson et al. 2011). Such dissimilarity coefficients provide complementary information regarding the main mechanisms responsible for β-diversity patterns (e.g., Siqueira et al. 2015).

In this study, we tested the hypothesis that ecological drift is a major process causing variation among small communities. To reach our goal, we used a unique, comprehensive dataset on aquatic insect communities sampled using identical methods in a total of 200 streams in two climatically highly different regions (100 in Brazil and 100 in Finland). The sampling design included 5 streams (communities) per watershed (20 in each region) and provided us replicates of metacommunities (watersheds; Fig. S1). This allowed us to make specific predictions considering a community size gradient. Our previous study showed that local community size is, on average, five-fold larger in boreal than in tropical streams, with the smallest boreal stream communities being as large as the largest tropical communities (Heino et al. 2018). Thus, we expected that β-diversity would be high (E1) and close (E2) to null expectations (random assembly from the regional pool) in watersheds with the smallest communities (some watersheds in Brazil only; Fig. 1). This would indicate that ecological drift plays a major role in structuring these small subtropical communities. Second, all else being equal, we expected that (E2a) β-deviation would be positive in all watersheds, but higher in the largest communities (Fig. 1). This would indicate that niche selection and sufficient dispersal rates are the main processes resulting in large communities to be more dissimilar than expected by chance, as species sorting occurs when dispersal is sufficient to allow individuals to reach sites that match their ecological requirements (Winegardner et al. 2012, Leibold and Chase 2018). Although dispersal limitation can also cause positive values of β-deviation, this was unlikely in our study system as the watersheds we studied are not large enough to lead to strong dispersal limitation (see details below). Taken together, these two expectations would lead to a negative relationship between β-diversity (before controlling for sampling effects) and community size, but to a positive relationship between β-deviation and community size (Fig. 1). Finally, because environmental heterogeneity and spatial extent are also expected to increase β-diversity (Heino et al. 2015a), we included them as co-variates in our models (Fig. S1).

**Figure 1.**
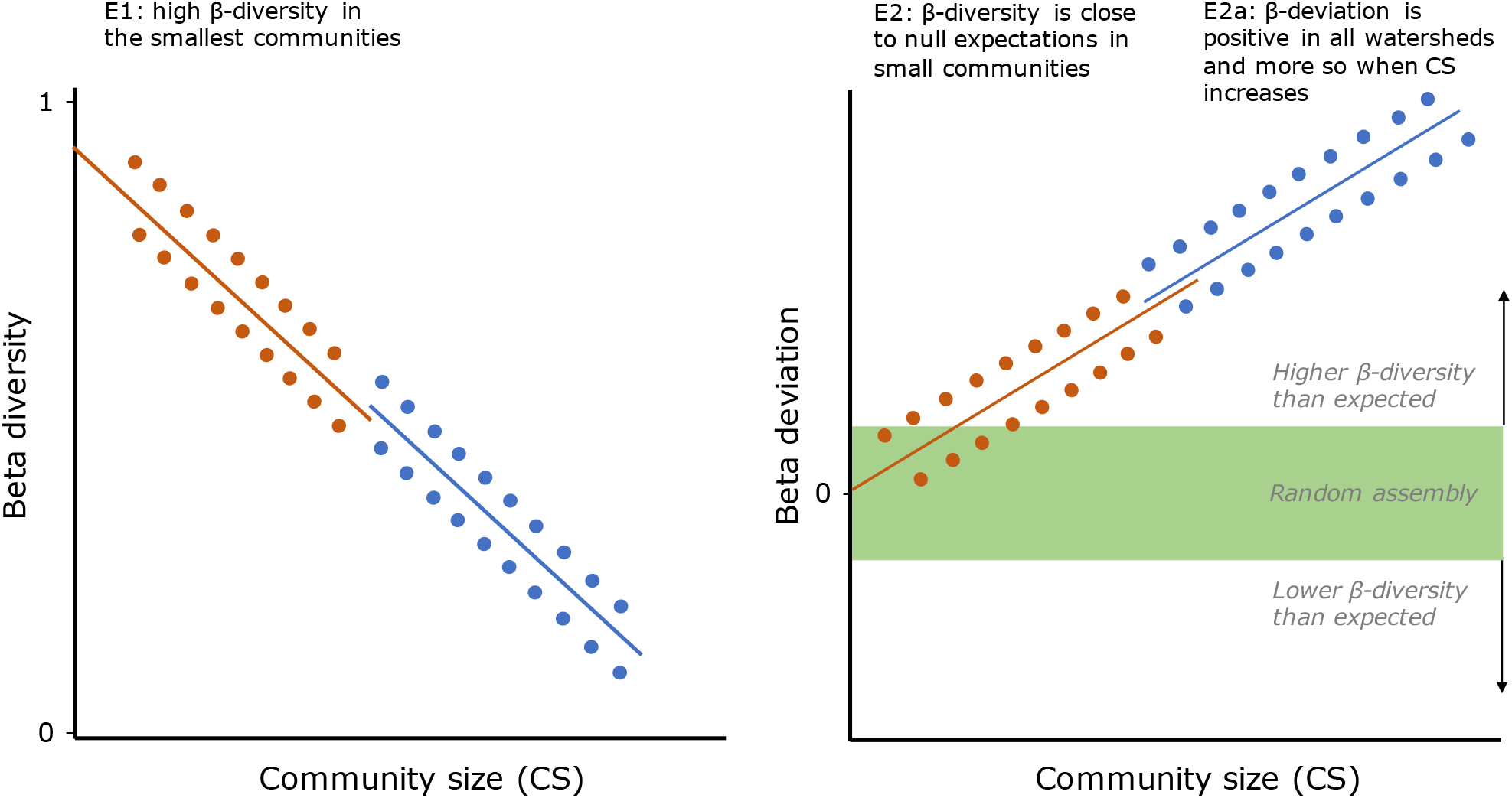
Graphical representation describing *a priori* expectations (E) about the relationship between beta diversity and beta deviation with community size in Brazil (vermilion) and Finland (blue).

## Material and methods

### Study area and sampling

The streams sampled in Brazil and Finland were distributed across a strong gradient of land cover among watersheds. It is well established that watershed land cover influences the structure of stream communities (Hynes 1975, Allan 2004, Roque et al. 2010, Siqueira et al. 2015). In Brazil, we sampled 100 streams distributed in 20 watersheds located in the southeastern region of the country – i.e., five streams per watershed, with spatial extent of 70 km in north-south and 120 km in east-west directions (Fig. S2). These streams drain through three major Atlantic Forest protected areas (‘Carlos Botelho’, ‘Intervales’ and ‘Alto Ribeira’ State Parks; São Paulo State). The watersheds are mainly dominated by pastures and planted forests (*Eucalyptus* and *Pinus*). The region has a dry season from April to August (average rainfall from 45 to 80 mm per month; average temperature from 16 to 20 °C) and a wet season from September to March (average rainfall from 105 mm to 180 mm per month; average temperature from 20 to 23 °C). Sampling was done between September and November in 2015.

The study sites in Finland were situated in the western part of the country. We sampled 100 streams that were distributed in 20 watersheds (as described above) with spatial extents of ca. 500 km and 300 km in north-south and in east-west directions (Fig. S2). The streams drain watersheds covered with agriculture and boreal forests. Western Finland has four typical seasons: a long winter that lasts from November to March, a short spring from April to May, a short summer from June to August, and an autumn period from September to October. Sampling was done in September 2014. Streams were generally of the same order within each watershed, but varied among watersheds, including 2^nd^ and 3^rd^ order streams in Brazil, and a few 4^th^ order streams in Finland. In Brazil, maximum distances between pairs of streams within watersheds varied from 2.48 to 8.86 km, whereas in Finland it varied from 12.77 to 109.5 km. Most in-stream abiotic variables varied within a similar range and had similar mean values between regions, except conductivity, total nitrogen and total phosphorus, which were much higher in Finland. A more detailed description of both regions can be found in Heino et al. (2018).

At each of the 100 stream sites in both regions, we took a 2-minute kick-net sample (net mesh size: 0.5 mm), which was composed of four 30-seconds sample units obtained in the main microhabitats at a riffle site (e.g., which considered differences in current velocity, depth, benthic particle size and macrophyte cover). The four sample units were pooled, preserved in alcohol in the field and taken to the laboratory for further processing and identification. All insects were separated from debris and the following taxonomic orders were identified to genus level: Ephemeroptera, Odonata, Plecoptera, Megaloptera, Trichoptera and Coleoptera. We sampled 16,113 individuals, distributed among 83 genera in Brazil, and 86,048 individuals (77 genera) in Finland. The mean number of genera per stream was 17.84 (standard deviation = 7.46) in Brazil and 14.01 (sd = 5.07) in Finland, while the mean number of individuals per stream was 181.50 (sd = 111.38) and 886.57 (sd = 700.73), respectively (Heino et al. 2018).

We adopted the definition of community size provided by Orrock and Watling (2010) and estimated local community size as the number of individuals sampled in a stream site. However, as β-diversity was estimated for each watershed (i.e., considering 5 stream sites; see below), we averaged the number of individuals across five streams within each watershed. This resulted in 20 values of community size, one per watershed (Fig. S1).

### Beta diversity and beta deviations

We first estimated β-diversity by using the Sørensen incidence-based coefficient and the Bray-Curtis abundance-based coefficient. To do that, we calculated dissimilarity values between all pairs of five streams within each of the 20 watersheds, separately for the tropical and boreal datasets. Our measure of β-diversity was represented by the mean of these values in each watershed. Thus, we estimated 20 values of Sørensen β-diversity and 20 values of Bray-Curtis β-diversity per region (Fig. S1).

We estimated β-deviations by using two procedures based on null models. To estimate incidence-based β-deviations that accounted for random sampling effects, we used a modified version of the Raup-Crick coefficient following Chase et al. (2011) and the procedures described in the package vegan (Oksanen et al. 2018): (i) we defined the genus pool as all genera occurring in each region and their observed occupancy across all 100 streams; (ii) an algorithm calculated the number of genera that every pair of streams share (*SG*_*obs*_); (iii) and assembled local stream communities by randomly sampling genus from the pool until reaching the local (observed) genus richness and by using the observed genus’ occupancy frequency to determine the probability to sample a genus; (iv) step (iii) was repeated 10,000 times to generate 10,000 random matrices of genus composition and, posteriorly, the number of genera shared between each pair of streams within each watershed (*SG*_*exp*_); (v) β-deviations were calculated as the number of random draws in which *SG*_*exp*_ ≥ *SG*_*obs*_ divided by the total number of random draws. This index was rescaled to range between −1 and 1, “indicating whether local communities are more dissimilar (approaching 1), as dissimilar (approaching 0), or less dissimilar (approaching −1), than expected by random chance” within each watershed (Chase et al. 2011). As with β-diversity, this procedure resulted in 20 values of incidence-based β-deviation per region, with one value per watershed (Fig. S1).

As selection and drift act on individuals, but not adding or removing entire populations, the use of abundance-based metrics should provide complementary information about assembly mechanisms. Thus, to estimate abundance-based β-deviations, we followed the procedure described by Kraft et al. (2011): (i) we defined the genus pool as all genera and their abundances across all 100 streams in each region; (ii) an algorithm then randomly assigned individuals to stream sites while preserving the overall genus-abundance distribution in the region and the total number of individuals in each stream site (local community size); (iii) step (ii) was repeated 10,000 to generate a distribution of pairwise dissimilarities within each watershed assuming random colonization of individuals from the genera pool; (iv) β-deviations were calculated as the difference between the observed Bray-Curtis dissimilarity and the mean expected (i.e. simulated) dissimilarity, divided by the standard deviation of the (null) simulated distribution within each watershed. Positive values and negative values indicate greater and lower dissimilarity than expected from random assembly, respectively. This null model allows one to analyze how β-diversity differs from patterns generated without the processes that cause clumping of species across the landscape (Kraft et al. 2011, Myers et al. 2015). These procedures also resulted in 20 values of abundance-based β-deviation per region, one value per watershed (Fig. S1).

We used linear regression models to test whether β-diversity (and β-deviation) was related to community size. Environmental heterogeneity within watersheds was estimated using environmental data and a Permutational analysis of multivariate dispersions (PERMDISP), as described by Anderson et al. (2006). This analysis was based on a Euclidean distance calculated with the standardized values (zero mean and unit variance) of the following variables: current velocity (m/s), depth (cm), stream width (cm), % of sand (0.25-2 mm), gravel (2-16 mm), pebble (16-64 mm), cobble (64-256 mm), and boulder (256-1024 mm), % of canopy cover by riparian vegetation, pH, conductivity, total nitrogen, and total phosphorus. These were all measured at the stream riffle scale (for details, see Heino et al. 2018). We also included the following watershed-scale variables estimated through satellite images within a 400-m buffer along tracts of the sampled streams: average slope, % of native forest cover, pasture, agriculture, planted forests, urban areas, mining, water bodies, bare soil, secondary forest cover, and mixed land uses. We mapped land use and land cover of Brazilian and Finnish watersheds using 5-m resolution RapidEye multispectral imagery and the CORINE database (https://land.copernicus.eu/pan-european/corine-land-cover), respectively. Geographical coordinates of the sampling sites were transformed to a Euclidean distance matrix and then submitted to a PERMDISP procedure (Anderson et al. 2006), using watershed as a grouping variable, to estimate mean spatial extent. Thus, we regressed our response variables (β-diversity and β-deviation; with 20 values; one per watershed) against community size, environmental heterogeneity, and spatial extent separately for each region (Fig. S1). We used the *vegan* package (Oksanen et al. 2018) in R (version 3.5.0; R Core Team 2018) to estimate β-diversity, Raup-Crick β-deviation, environmental heterogeneity, and spatial extent. In the tropical data, the strongest correlation between explanatory variables was between environmental heterogeneity and spatial extent (r = 0.47), whereas in the boreal data, it was between community size and spatial extent (r = 0.36). All other correlations were lower than 0.07. Thus, multicollinearity was not an issue in our study.

## Results

There was a strong negative relationship between β-diversity (both with incidence- and abundance-based data) and community size in Brazil (*b* = −0.87, *t* = −7.47, *p* < 0.001, *R*^2^ = 0.74; and *b* = −0.78, *t* = −5.32, *p* < 0.001, *R*^2^ = 0.59, respectively; Fig. 2), indicating that small communities were more dissimilar among each other than larger communities, as we expected (E1). However, we found that neither incidence-based (Sørensen) nor abundance-based (Bray-Curtis) β-diversity were related to community size in Finland (*b* = −0.14, *t* = −0.59, *p* = 0.563; and *b* = −0.22, *t* = −0.94, *p* = 0.357, respectively).

**Figure 2.**
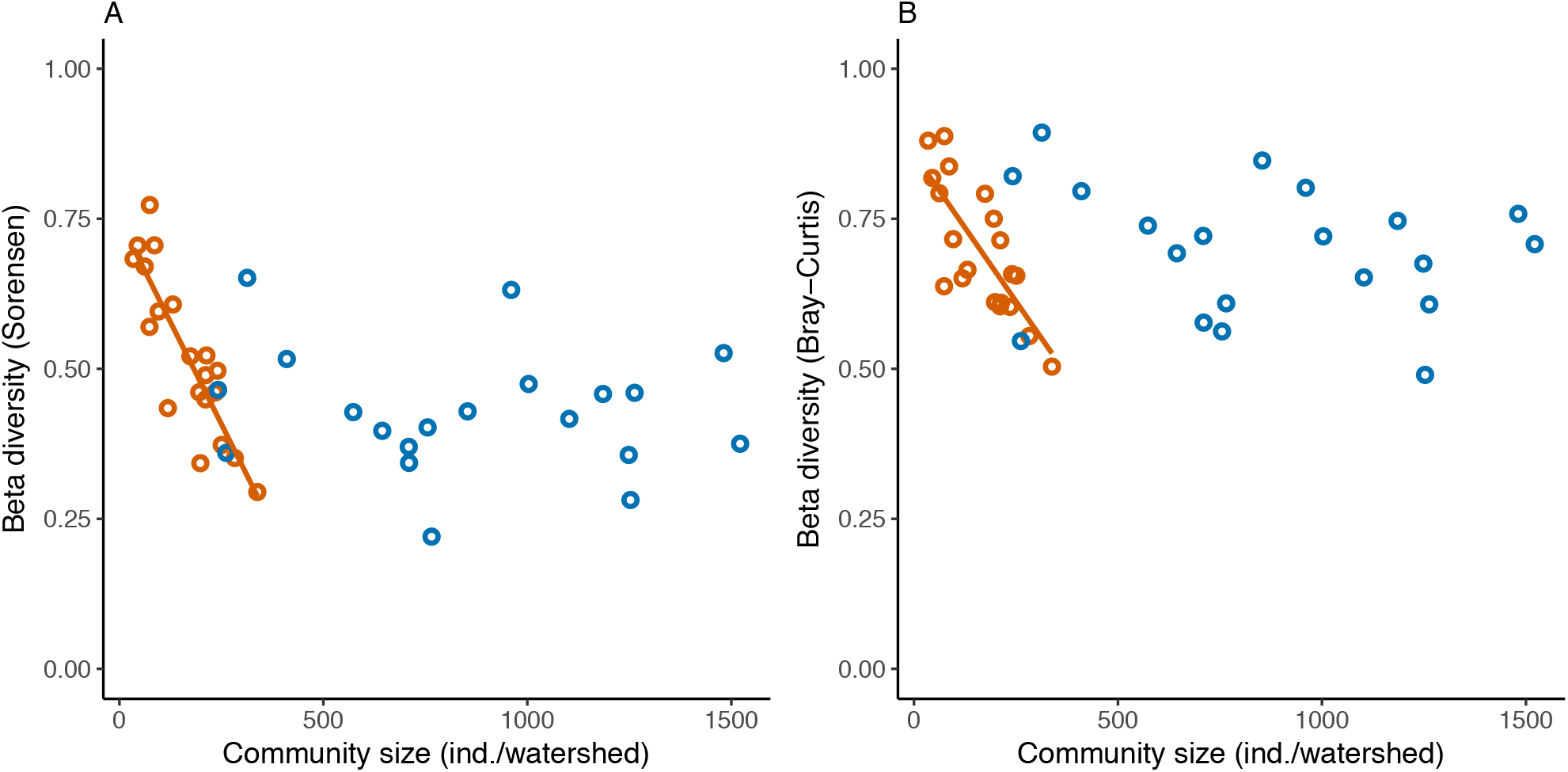
(A) Incidence-based (Sørensen) and (B) abundance-based (Bray-Curtis) beta diversity-community size (average number of individuals per watershed) relationships within tropical (vermilion) and boreal (blue) stream watersheds (*n* = 20 for each region). The average number of individuals per watershed was calculated with a sample size of 5 streams.

We found strong relationships between β-deviation (i.e., β-diversity apart from null expectations) and community size in both regions, but, contrary to our expectations, they varied according to the type of dissimilarity coefficient (incidence- and abundance-based) and the direction of relationship (negative and positive; Table 1, Fig. 3). As we expected (E2), beta diversity of tropical smaller communities was closer to null expectations than those of larger communities (Fig. 3). However, differently from what we expected, mean Raup-Crick β-deviations were negative in Brazil, varying from −0.98 to −0.02, indicating that tropical communities were less dissimilar in genus composition than expected by random sampling from the species pool. Neither environmental heterogeneity nor spatial extent were related to Raup-Crick β-deviation in Brazil. Raup-Crick β-deviation in boreal watersheds (ranging from −0.99 to 0.30) was not related to community size (Fig. 3A) but it was to environmental heterogeneity (Table 1).

**Table 1.** Relationship between beta deviation (incidence-based [Raup-Crick] and abundance-based [Bray-Curtis]) and community size, environmental heterogeneity and spatial extent (*n* = 20 watersheds in each region). CS = Community size; EH = Environmental heterogeneity; SE = Spatial extent. *R*^2^ and adj. *R*^2^ = coefficient of determination and adjusted coefficient of determination of the full model, respectively. *b* = standardized regression coefficient.

| | | | $b$ | $se$ | $t$ | $P$ | $R^2$ | adj. $R^2$ |
| --- | --- | --- | --- | --- | --- | --- | --- | --- |
| Brazil | Raup-Crick | CS | **-0.737** | **0.164** | **-4.505** | **<0.001** | 0.510 | 0.419 |
|  |  | EH | -0.114 | 0.200 | -0.572 | 0.575 |  |  |
|  |  | SE | 0.057 | 0.200 | 0.289 | 0.776 |  |  |
|  | Bray-Curtis | CS | **0.777** | **0.147** | **5.279** | **<0.001** | 0.658 | 0.594 |
|  |  | EH | 0.211 | 0.167 | 1.267 | 0.223 |  |  |
|  |  | SE | -0.087 | 0.167 | -0.522 | 0.609 |  |  |
| Finland | Raup-Crick | CS | -0.252 | 0.221 | -1.137 | 0.271 | 0.319 | 0.192 |
|  |  | EH | **0.461** | **0.207** | **2.229** | **0.040** |  |  |
|  |  | SE | 0.338 | 0.222 | 1.527 | 0.146 |  |  |
|  | Bray-Curtis | CS | **0.728** | **0.168** | **4.328** | **<0.001** | 0.607 | 0.534 |
|  |  | EH | -0.115 | 0.157 | -0.731 | 0.475 |  |  |
|  |  | SE | 0.087 | 0.168 | 0.518 | 0.611 |  |  |

**Figure 3.**
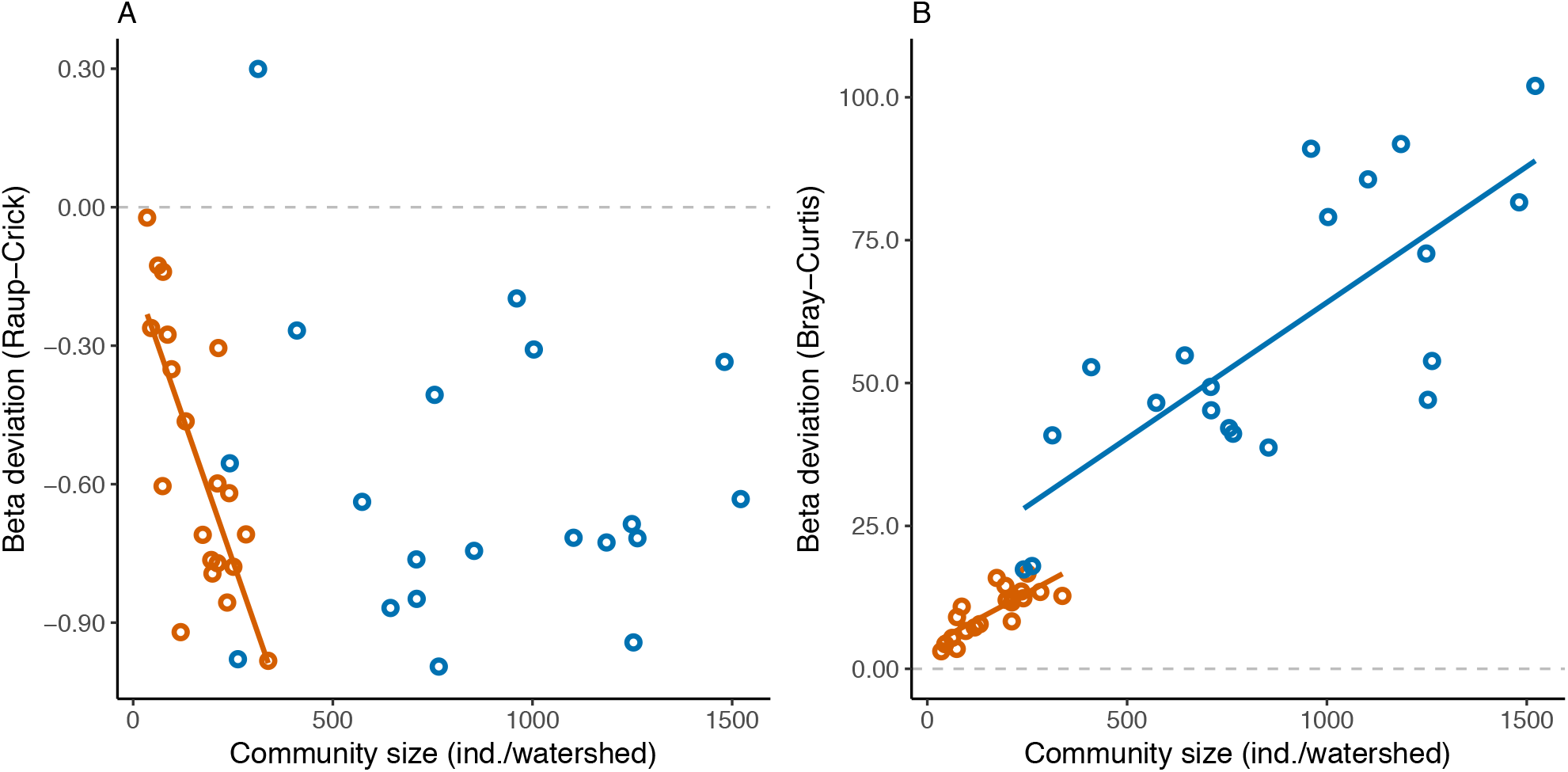
(A) Incidence-based (Raup-Crick) and (B) abundance-based (Bray-Curtis) beta deviation-community size (average number of individuals per watershed) relationships within tropical (vermilion) and boreal (blue) stream watersheds (*n* = 20 for each region). The average number of individuals per watershed was calculated with a sample size of 5 streams. The dashed grey line indicates expected beta diversity under null assembly.

Bray-Curtis (abundance-based) β-deviation was positively related to community size in both regions (Table 1; Fig. 3B), supporting our expectations (E2a). In tropical watersheds, mean Bray-Curtis β-deviations ranged from 3.08 to 16.66, indicating that communities were more dissimilar than expected by random changes in species abundances and genus composition. Again, β-deviations of smaller communities were closer to zero than those of larger tropical communities, supporting our expectation E1. In boreal watersheds, mean Bray-Curtis β-deviation varied from 17.40 to 102.51, also indicating that communities were more dissimilar than expected by random changes in genus abundances and composition. Neither environmental heterogeneity nor spatial extent were related to Bray-Curtis β-deviation in any region.

Although there was a negative relationship between Raup-Crick β-deviation and community size in tropical watersheds only, mean incidence-based β-deviation in Brazil was not different from mean incidence-based β-deviation in Finland (Fig. 3A). This suggests that the departure from null expectations related to compositional change among streams is similar in tropical and boreal regions. On the other hand, mean abundance-based β-deviation was five times higher in Finland than in Brazil, suggesting that departures from null expectations are much higher in boreal than in tropical streams (Fig. 3B).

## Discussion

The role of stochastic community assembly processes in arranging species within and among communities has gained support from models (Mouquet and Loreau 2003, Orrock and Fletcher 2005, Durães et al. 2016) and data (Cottenie 2005, Lancaster and Downes 2017, Germain et al. 2017, Swan and Brown 2017, Valente-Neto et al. 2017). Here, we provide empirical evidence that community size, a simple characteristic of communities, may mediate the interplay between niche selection and ecological drift as drivers of β-diversity in tropical and boreal metacommunities. Based on null models, our estimates of variation in community structure accounted for both differences in species richness and species relative abundance that deviate from random assembly (i.e., β-deviations). We showed that β-diversity of smaller tropical communities deviates less from null expectations than larger communities. This means that the high β-diversity we observed among smaller communities was, to some extent, indistinguishable from patterns generated by random changes in local genus richness and community size. As findings generated by null models are our best approximation of patterns generated by stochastic processes (Kraft et al. 2011, Chase et al. 2011), we suggest that demographic stochasticity plays a major role in small ecological communities (Orrock and Fletcher 2005, Orrock and Watling 2010, Gilbert and Levine 2017).

Explanations for the major role of demographic stochasticity in small communities involve the alteration of the occupancy frequency and relative abundance of species with different fitness (Orrock and Watling 2010). When local communities are small, even species with high fitness are at a high risk of extinction due to demographic stochasticity in comparison to a situation when communities have large populations. Consequently, species with low relative abundances have a chance to increase their populations in small communities (Orrock and Watling 2010, Gilbert and Levine 2017). As this would be the result of a local demographic random process, and because dispersal ability is highly variable among species in riverine systems (Heino et al. 2015b, Tonkin et al. 2018a), the outcome of the assembly would likely differ among local communities, increasing β-diversity within metacommunities. Indeed, in Brazil, the three most abundant and widespread genera (Heino et al. 2018) were not dominant in smaller communities: the mayfly *Farrodes* was not among the most abundant in any of the smallest communities; the beetle *Heterelmis* was among the most abundant in two communities only; and the caddisfly *Smicridea* was among the most abundant in three of the five smallest communities. These three genera were dominant in four of the five largest communities in Brazil. The smallest communities in Brazil were dominated by genera with intermediate regional abundance and occupancy. For example, *Gripopteryx* (stonefly), *Cloeodes* (mayfly) and *Callibaetis* (mayfly) were the sixth, eleventh and sixteenth most abundant genera in Brazil, respectively. In Finland, these differences were less evident, as the three most abundant and widespread genera dominated small and large communities: the beetle *Elmis* (two of the smallest vs. three of the largest communities); the mayfly *Baetis* (three vs. three); and stonefly *Nemoura* (one vs. one). This is likely because the smallest boreal stream communities were as large as the largest tropical communities (Heino et al. 2018).

The positive relationship between abundance-based β-deviation and community size in both regions is in line with our expectations. The positive slope and values of β-deviation indicate that the variation in community structure among streams increased with community size more than what was expected under random assembly. Thus, in terms of which genera were more abundant or less abundant, and more aggregated or less aggregated, communities within the same watershed differed from each other more than expected by chance, especially in mid to large communities. As dispersal within watersheds was likely not limited, this positive relationship indicates that niche selection was sufficient to cause non-random variations in genera relative abundance and aggregation patterns among large communities. We suggest that as community size increases, demographic stochasticity becomes less important and niche selection determines which species are more abundant locally and widely distributed within the metacommunity. In this case, small random variations in the number of individuals of relatively abundant genera occurring in larger communities, which can only be detected with abundance-based β-diversity metrics, do not result in major changes in genus occurrence.

Although small communities had incidence-based β-deviations values close to random assembly expectations, as we expected, the relationship between β-deviation and community size was negative in Brazil. This result contradicts our expectations and indicates that the genus composition of streams harboring mid to large communities in tropical watersheds is less dissimilar (negative values of incidence-based β-deviation) than patterns predicted by random assembly – i.e., these communities share more genera than expected. In general, communities can be less dissimilar than random expectations in at least two cases. First, dissimilarity should be low when niche selection is spatially constant, as the environment maximizes the fitness of a few species (Vellend 2016). Within watershed environmental heterogeneity was weakly and positively related with incidence-based β-deviation in Finland only, where community sizes are on average five-fold larger than in tropical streams (Heino et al. 2018). This suggests a tendency towards environmental determinism; the higher the environmental variation within watersheds, the higher their β-deviation. Thus, it is likely that even the large size of boreal communities was not sufficient to allow niche selection to be the main driver of spatial variation in genus composition among those communities. Alternatively, we might have failed to measure key abiotic and biotic variables (e.g., biological interactions) underlying community variation, or despite including key correlates of beta diversity, our snapshot analysis might not have been enough to represent the complexity of all mechanisms involved in niche selection in watersheds.

A second reason for communities to be less dissimilar than random expectations is when dispersal rates are high within the metacommunity (Mouquet and Loreau 2003, Leibold and Chase 2018). By distributing organisms among communities, dispersal can reduce β-diversity if it is excessive and is combined with a source-sink system, where populations with high growth rates supply individuals to other localities where they would otherwise be excluded by niche selection (Mouquet and Loreau 2003, Siqueira et al. 2014). Although dispersal across watersheds tends to be limited for many aquatic species, those with an adult flight stage may have higher dispersal rates especially along the stream channel (Hughes 2007, Lancaster and Downes 2017). A meta-analysis by Muehlbauer et al. (2014) showed that most adult aquatic insects tend to fly ca. 1.5 m around their natal stream, but that a few individuals can fly 550 m away from the stream, with some caddisflies being able to reach sites distant more than 650 m. Also, while Macneale et al. (2005) found that the stonefly *Leuctra ferruginea* was able to fly across headwater forested catchments, Flenner and Sahlin (2008) estimated annual range expansions of up to 88 km in non-migratory dragonflies in a boreal region of Sweden. The northern faunas of Europe should also be composed of species with good dispersal ability, as they have reached these areas since the Last Glacial Period. On the other hand, many tropical aquatic insects have multiple reproduction events per year, i.e., many are multivoltine species (Wallace and Anderson 1996, Vásquez et al. 2009). Unless local population growth rate is low, multivoltine species should have many opportunities per year to disperse from their natal streams. If a few of these multivoltine species can fly 250 m (a conservative estimate considering the results by Muehlbauer et al. 2014), after some generations, some individuals of these species could reach distant sites, at a velocity of one kilometer per year. We thus suggest that long-term availability for colonization and multivoltinism allowed some genera to reach widespread distribution within tropical watersheds (Saito et al. 2016). Similarly, good dispersal abilities of northern species may have masked the role of within-watershed environmental heterogeneity in the boreal region, making these communities less dissimilar than expected. Thus, as communities become larger, demographic stochasticity plays a less important role, while dispersal surplus plays a major role in homogenizing the genus composition of streams within tropical watersheds. However, this, as almost everything in ecology, should be scale dependent (Chase et al. 2018). If our watersheds were larger (in extent), dispersal limitation would likely have played a role and niche selection could have been the major driver of community structure. Our results reinforce previous suggestions that dispersal should be viewed as a process occurring over time and not only considering the distance individuals can move in one dispersal event during one generation (Saito et al. 2015).

Predicting community assembly dynamics has been a major challenge for ecology in a changing world (Mouquet et al. 2015). Many drivers of environmental change, such as habitat destruction, overexploitation, pollution, and reductions in landscape connectivity, are likely to cause reductions in community size. For example, Hallmann et al. (2017) estimated a decline of more than 70% in insect biomass between 1989 and 2016 in Germany and Lister and Garcia (2018) found that arthropod biomass has fallen 10 to 60 times since 1970 in Puerto Rico. It is possible, then, that results from previous studies associating high β-diversity with environmental changes were related to small communities being more variable. For example, Hawkins et al. (2015) showed that disturbed sites had high β-diversity due to a decreased prevalence of more common taxa. Although they did not analyze community size per se, the mechanism they evoked to explain this pattern – i.e., the relative abundance of species with lower fitness become progressively higher – is what has been suggested as the reason for a major role of ecological drift in small communities (Orrock and Watling 2010, Gilbert and Levine 2017). Additionally, the effects of environmental changes on naturally small communities might be combined with the effects of ecological drift in non-obvious ways, making community dynamics even harder to predict (e.g., Bini et al. 2014) and smaller communities more vulnerable to novel environmental conditions, such as altered flow regimes (Tonkin et al. 2018b, Ruhi et al. 2018). This should also be relevant for the conservation and management of ecosystems. For example, stream restoration efforts tend to be directed towards sites with low habitat quality and with reduced number of species and low-density populations. Even if restoration efforts overcome dispersal constraints and barriers to recolonization, which are a major cause for unsuccessful stream restoration (Bond and Lake 2003, Sundermann et al. 2011, Tonkin et al. 2014), restored communities will likely be small at the earlier stages of community assembly and, thus, more prone to ecological drift.

Our study provides empirical evidence of the role that community size can play in mediating stochastic and deterministic process as drivers of metacommunity dynamics in tropical and boreal streams, providing one possible solution to a long-standing debate about why some communities are apparently more influenced by stochastic processes than others. The magnitude of β-diversity deviation from the null models (negative and positive values) and how it related with community size (negative and positive slopes) indicate that stochastic and deterministic processes affect species occurrences and their abundances differently. While incidence-based β-deviation was negative and decreased with community size in Brazil, abundance-based β-deviation was positive and increased with community size in both regions. These results indicate that: (1) as communities get larger, demographic stochasticity plays a less important role and excessive dispersal combined with niche selection tend to homogenize the genus composition of larger communities; and (2) niche selection was likely strong enough to produce variations in genera relative abundances, but not to cause compositional changes among communities within the same watershed. However, we acknowledge that these are, at least partially, *a posteriori* explanations as we expected that the results would be similar (i.e., a positive relationship between beta deviation and community size) independently of the type of coefficient. On the other hand, these contrasting results appear interesting avenues for future research. Our findings complement previous empirical and conceptual efforts that indicated that ecological drift can drive biodiversity patterns even in communities where species have clear differences in life-history traits, resource use and competitive abilities (i.e., those communities assumed to be non-neutral). Incorporation of community size into ecological models should provide conceptual, empirical and applied insights towards better understanding of the processes driving changes in biodiversity.

## Data accessibility

Code and data that fully reproduce the analyses are archived on Zenodo (https://doi.org/10.5281/zenodo.2620550).

## Acknowledgements

This preprint has been peer-reviewed and recommended by Peer Community In Ecology (https://dx.doi.org/10.24072/pci.ecology.100023). We thank Eric Harvey, Kevin Cazelles, and Romain Bertrand for their comments and suggestions on earlier versions of this manuscript. This research originated during a joint research project funded by grant #2013/50424-1 (São Paulo Research Foundation, FAPESP) to TS, and by grants #273557 and #273560 to JH and JS (Academy of Finland). ASM and LMB are supported by research grants from the Conselho Nacional de Desenvolvimento Científico e Tecnológico (CNPq no. 307587/2017-7 and 304314/2014-5, respectively). This study was financed in part by the Coordenação de Aperfeiçoamento de Pessoal de Nível Superior – Brasil (CAPES) – Finance Code 001 (scholarship to DKP). This work was also developed in the context of the National Institutes for Science and Technology (INCT) in Ecology, Evolution and Biodiversity Conservation, supported by MCTIC/CNPq (proc. 465610/2014-5) and FAPEG. We thank Christopher Catano and Jonathan Myers for gently providing R code to generate abundance-based beta deviations, and Mauricio Vancine for preparing a map of the regions.

## Author contributions

TS conceived the idea, ran the analyses, and wrote the manuscript with substantial support from VSS, LMB, ASM, and JH. TS, LMB, VLL, ASM, JS, and JH planned the sampling design. DKP and JJ-M sampled the data. DKP, VSS and KTT identified the insects. DKP, JJ-M, KT, VLL and JS commented on the manuscript.

## Supplementary material

**Figure S1.**
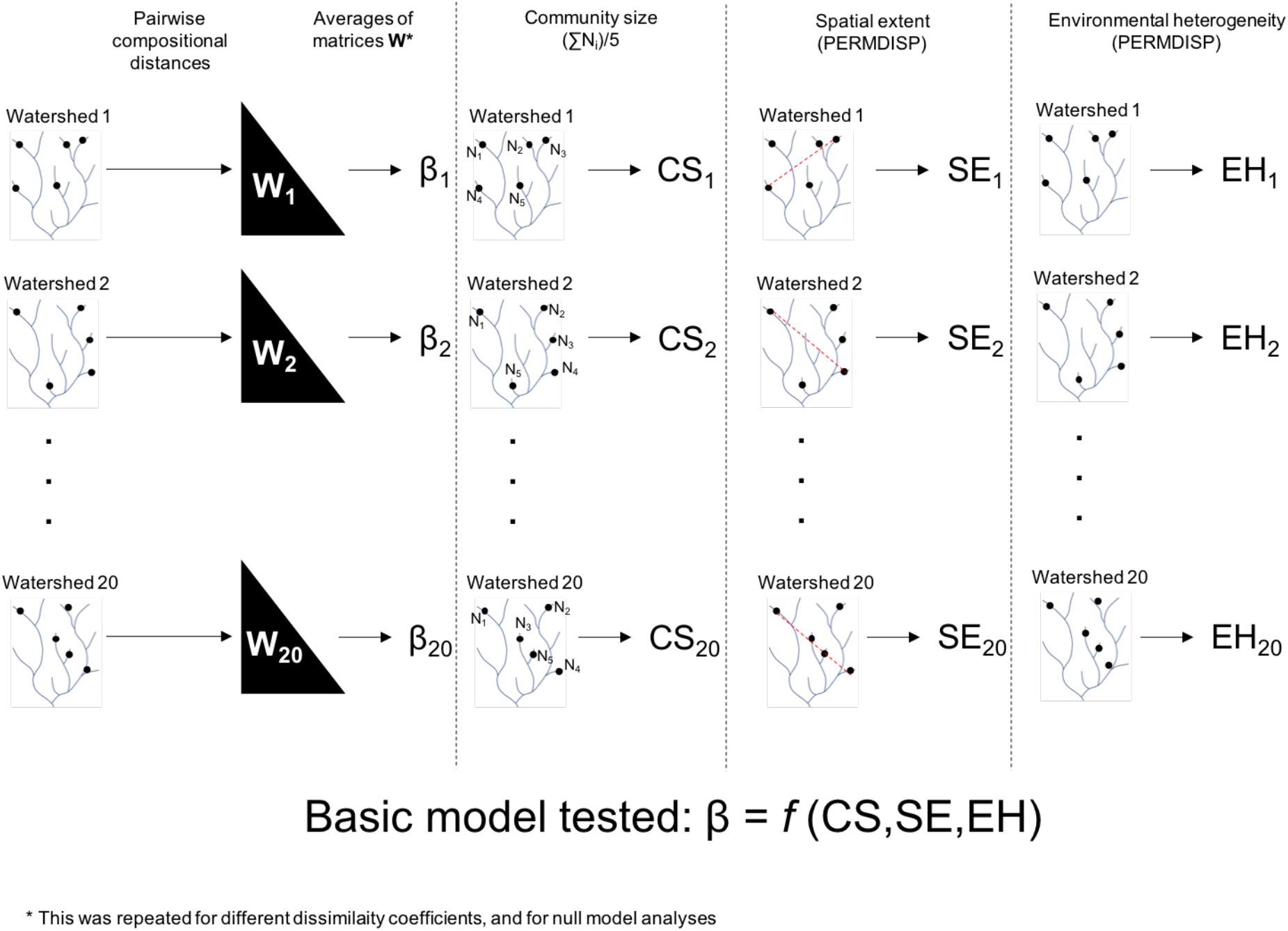
Graphical representation describing the general procedures for estimating beta diversity (β), community size (CS), spatial extend (SE) and environmental heterogeneity (EH) within watersheds (*n* = 20) in Brazil and Finland.

**Figure S2.**
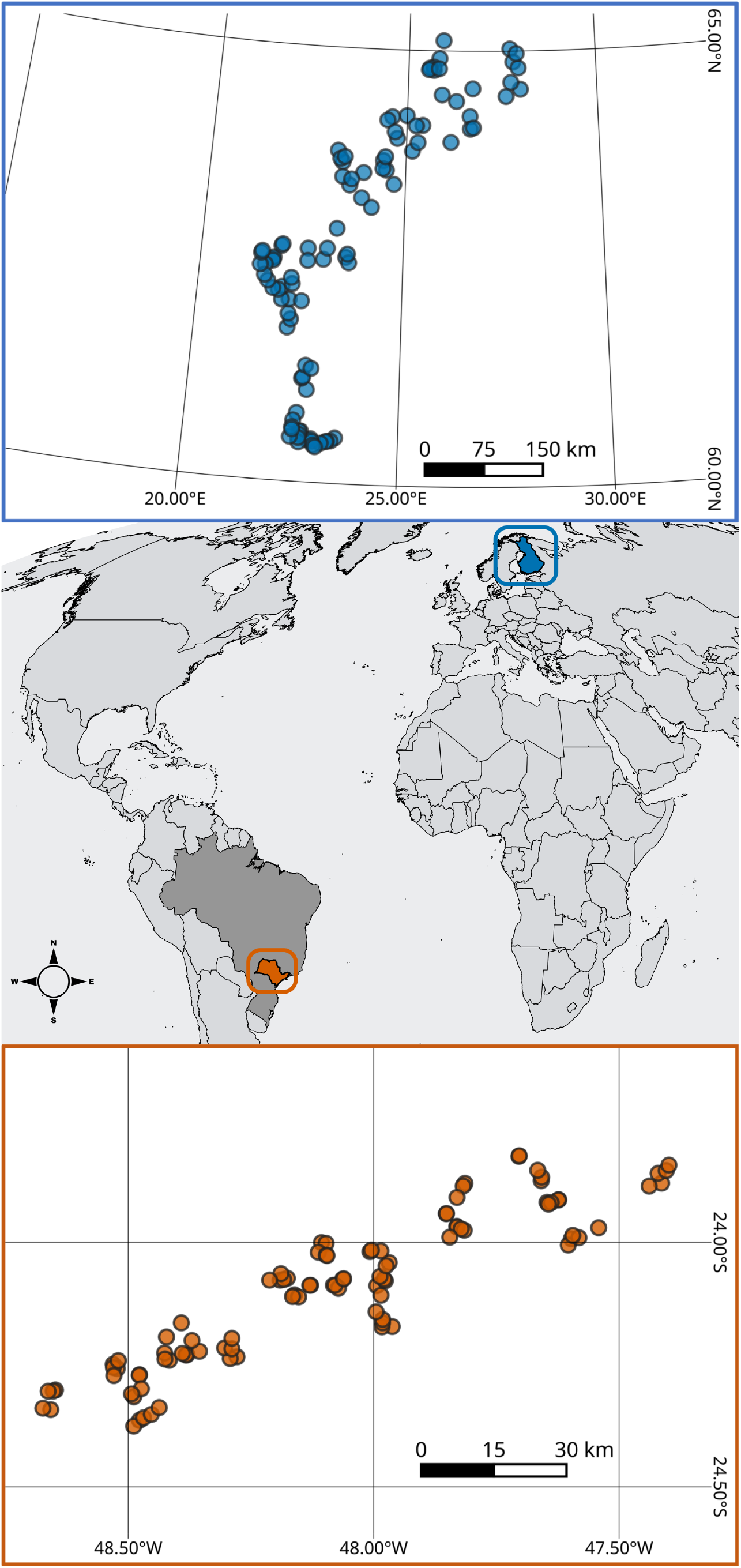
Geographical location and spatial distribution of streams in São Paulo State, Brazil (circles in vermilion) and Finland (circles in blue).

